# ASK1 overexpression protects from HFD-induced body weight gain via FGF21

**DOI:** 10.1101/2025.01.23.634369

**Authors:** Anne Goergen, Tenagne D. Challa, Marcela Borsigova, Carlos Villarroel-Vicente, Pim van Krieken, Stephan Wueest, Daniel Konrad

## Abstract

People with obesity are at high risk to develop metabolic complications such as type 2 diabetes and metabolic dysfunction-associated fatty liver disease (MAFLD). We previously reported that the apoptosis signal-regulating kinase 1 (ASK1) regulates the development of MAFLD, since high fat diet (HFD)-fed mice with liver-specific ASK1 depletion and overexpression revealed increased and blunted development of MAFLD, respectively. Herein we identify a protective role of liver-expressed ASK1 in the context of diet-induced obesity. When fed a HFD for 20 weeks, liver-specific ASK1 overexpressing mice (ASK1^+hep^) were resistant to obesity and showed improved glucose metabolism as well as increased energy expenditure compared to control animals. Moreover, HFD-fed ASK1^+hep^ mice had higher BAT activity as well as increased browning of inguinal WAT, as suggested by increased UCP1 levels. The latter may be induced by the hepatokine fibroblast growth factor (FGF21), a hormone that is mainly produced in the liver and is known to reduce body weight via increasing energy expenditure. In line with an important role of FGF21, its plasma levels were significantly increased in HFD-fed ASK1^+hep^ mice and they negatively correlated with body weight. Mechanistically, we propose that the transcription factor ATF4 is activated via HFD-induced ASK1-p38 signaling in hepatocytes, subsequently promoting hepatic *Fgf21* gene expression. Supporting this hypothesis, p38 inhibition resulted in a more pronounced reduction in *Fgf21* expression in ASK1^+hep^ hepatocytes compared to controls, while silencing of ATF4 in primary hepatocytes significantly decreased *Fgf21* transcript levels in ASK1^+hep^ hepatocytes but not in hepatocytes derived from control mice. In conclusion, we have uncovered a yet undescribed axis between hepatic ASK1 and FGF21 which positively affects body weight and glucose metabolism.

## 1. Introduction

The rising prevalence of overweight and obesity exhibits a heavy burden for societies and health-care systems all over the world (1, 2). Obesity is often paralleled by the occurrence of metabolic comorbidities such as cardiovascular incidents, type 2 diabetes and metabolic dysfunction-associated fatty liver disease (MAFLD) (3). Importantly, cardiovascular and metabolic diseases are the primary causes of death and disability globally, impacting the quality of life for millions of people (4, 5).

Fibroblast growth factor 21 (FGF21) is a stress-induced hormone, mainly secreted from the liver, that has fundamental functions in the regulation of glucose and energy metabolism (6). Interest in the metabolic, and more importantly therapeutic effects of FGF21 was first stimulated by the finding that FGF21 induces insulin-independent glucose uptake in murine adipocytes. Moreover, FGF21 protected mice from diet-induced obesity and lowered blood glucose concentrations in diabetic mice (7–9). One mechanism by which these effects occur is suggested to be via an increase in thermogenesis-driven energy expenditure by FGF21 (10). Indeed, it has repeatedly been shown that FGF21 leads to enhanced energy expenditure via the activation of brown adipose tissue (BAT) and induction of inguinal white adipose tissue browning (ingWAT) (11–14). Thermogenesis is mainly, but not exclusively, driven by uncoupling protein 1 (UCP1), a protein that uncouples the proton gradient over the inner mitochondrial membrane from ATP synthesis, thus creating a futile cycle that produces heat (15). Endocrine FGF21 analogs have thus gained attention for their potential to directly target the adipose tissue and the liver, thereby promoting a healthier state of whole-body metabolism (8, 9, 16, 17). However, pharmacological studies have also revealed adverse effects of administering supra-physiological doses of exogenous FGF21 (18, 19), underlining the need to find alternative methods of elevating circulating FGF21 *in vivo*.

MAFLD and its progression to metabolic dysfunction-associated hepatic steatohepatitis (MASH) as well as liver fibrosis are important comorbidities of the current obesity pandemic and constitute the most frequent liver diseases world-wide (20, 21). We previously reported that the apoptosis signal-regulating kinase 1 (ASK1) blunts the development of MAFLD and liver fibrosis. In particular, we demonstrated that hepatic ASK1 knockout impaired glucose metabolism and accelerated the development of fatty liver disease, hepatic inflammation and liver fibrosis in high fat diet (HFD)-fed mice. Conversely, liver-specific ASK1 overexpressing mice (ASK1^+hep^) were protected from HFD-induced hepatic lipid deposition, suggesting a protective role of liver-expressed ASK1 in MAFLD (22). Moreover, we observed that ASK1^+hep^ mice were resistant to HFD-induced obesity and associated metabolic derailments such as impaired glucose tolerance. Thus, in the present study, we aimed to understand how liver—specific overexpression of ASK1 confers resistance to body weight gain upon HFD feeding in mice. We hypothesize that ASK1 induces the expression of a hepatokine that either decreases food intake or increases energy expenditure and, thereby, decreases body weight gain.

## 2. Results

### ASK1^+hep^ mice are resistant to HFD-induced obesity

We have previously shown that liver-specific ASK1 overexpressing (ASK1^+hep^) mice are partly protected from HFD-induced hepatic steatosis (22). To elucidate the role of liver-specific ASK1 overexpression beyond its protective role in the development of MAFLD, we studied the effect of long-term HFD feeding on glucose and energy metabolism in ASK1^+hep^ mice. To this end, ASK1^+hep^ and control littermate (ASK1^f/f^) mice were fed a regular chow or HFD for 20 weeks. Upon HFD feeding, ASK1^+hep^ mice gained significantly less body weight compared to littermate controls (Fig. 1A). ASK1^+hep^ mice were on average ∼10 grams lighter than littermate control mice after the 20-week HFD challenge, which was mirrored by significantly lower total white adipose tissue and liver weights (Fig. 1B), implying that ASK1^+hep^ mice are resistant to diet-induced obesity. In parallel, HFD-fed ASK1^+hep^ mice revealed significantly improved glucose tolerance (Fig. 1C) as well as ameliorated insulin sensitivity (Fig. 1D) compared to ASK1^f/f^ mice. Consistently, HFD-fed ASK1^+hep^ mice showed lower fasting blood insulin and glucose concentrations compared to littermate controls (Suppl. Fig. 1A, B). While 20-week HFD-fed ASK1^f/f^ mice revealed significantly increased body weight compared to chow-fed mice as expected (HFD-fed ASK1^f/f^ 46.7±1.8g *vs.* chow-fed ASK1^f/f^ 38.5±0.8g, p<0.001), chow-fed ASK1^f/f^ and ASK1^+hep^ mice did not significantly differ in body weight, glucose tolerance and insulin sensitivity at the age of 26 weeks (Suppl. Fig. 1 C-E). To summarize, liver-specific ASK1 overexpression protects mice from HFD-induced body weight gain and associated deterioration in glucose tolerance and insulin sensitivity.

**Figure 1.**
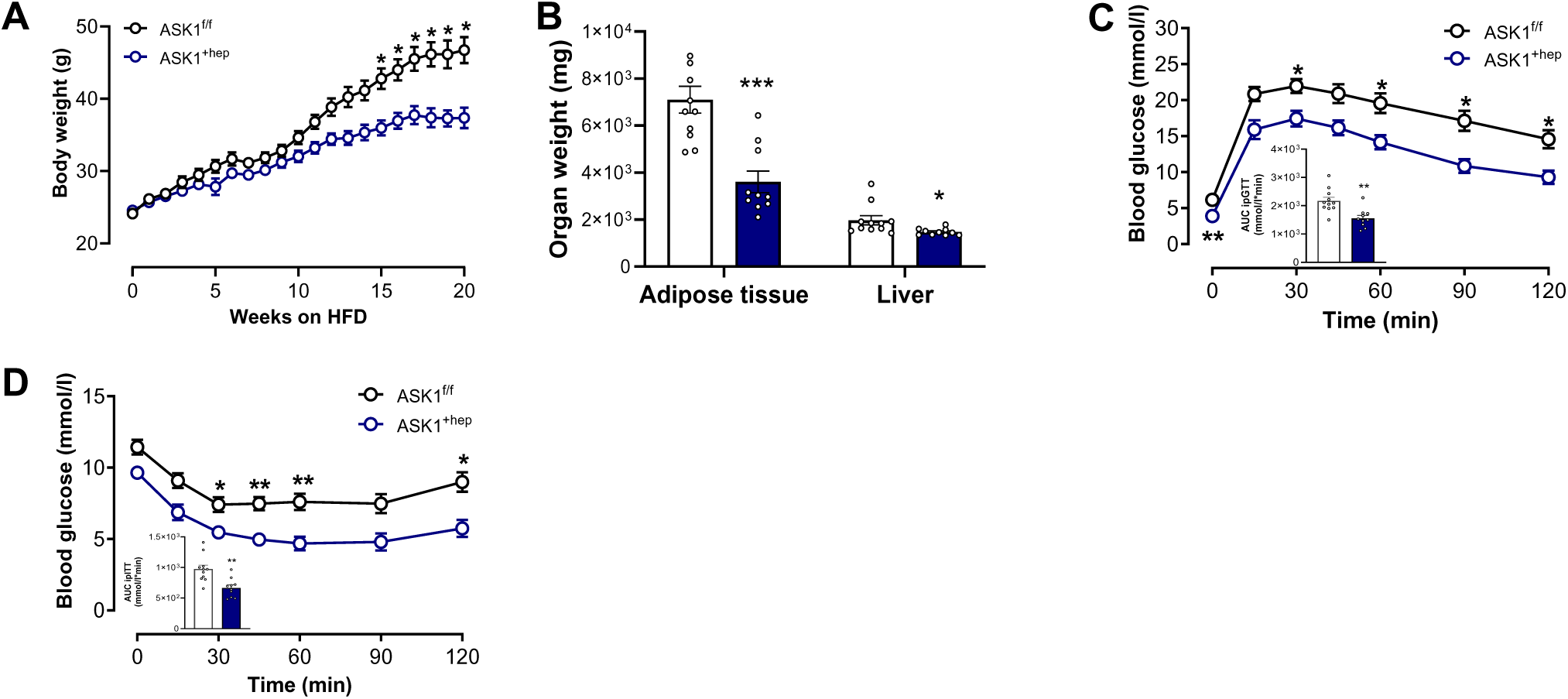
ASK1^+hep^ mice are resistant to HFD-induced obesity. **A**. Body weight development (grams) over time (weeks) in HFD-fed ASK1^f/f^ and ASK1^+hep^ mice (n=10-11 mice). **B.** Organ weight (mg) of 20 weeks HFD-fed ASK1^+hep^ (blue bar) and ASK1^f/f^ (white bar) (n=10-12 mice). **C.** Intraperitoneal glucose tolerance test (ipGTT) including area under curve (AUC) (**C**) and insulin tolerance test (ipITT) including area under curve (AUC) (**D**) in ASK1^+hep^ and ASK1^f/f^ mice fed a HFD for 20 weeks (n=10-11 mice). Data are shown as mean ± SEM. *p<0.05, **p<0.01, ***p<0.001 (Two-way ANOVA for panels A, C and D, t-test for panel B).

### Energy expenditure is increased in HFD-fed ASK1^+hep^ mice

Next, we sought to understand why ASK1^+hep^ mice are protected from HFD-induced body weight gain. To this end, we analyzed whole body physiology and molecular composition of metabolic tissue in mice after 7 weeks of HFD (Fig. 2A), a time point when body weight started to deviate between the two genotypes (Fig. 1A). From a thermodynamic point of view, reduced body weight may originate from decreased food intake or increased energy expenditure. Thus, these parameters were assessed in metabolic cage units. Linear regression analysis revealed a significant body weight-independent increase in energy expenditure in HFD-fed ASK1^+hep^ mice (Fig. 2B), while food intake was not different between the genotypes (Fig. 2C), indicating that reduced body weight gain in HFD-fed ASK1^+hep^ mice originates from increased energy expenditure. Of note, locomotor activity was not increased in HFD-fed ASK1^+hep^ compared to ASK1^f/f^ mice (Suppl. Fig 1F), suggesting that increased thermogenesis rather than elevated locomotion increased energy expenditure in overexpressing mice.

**Figure 2.**
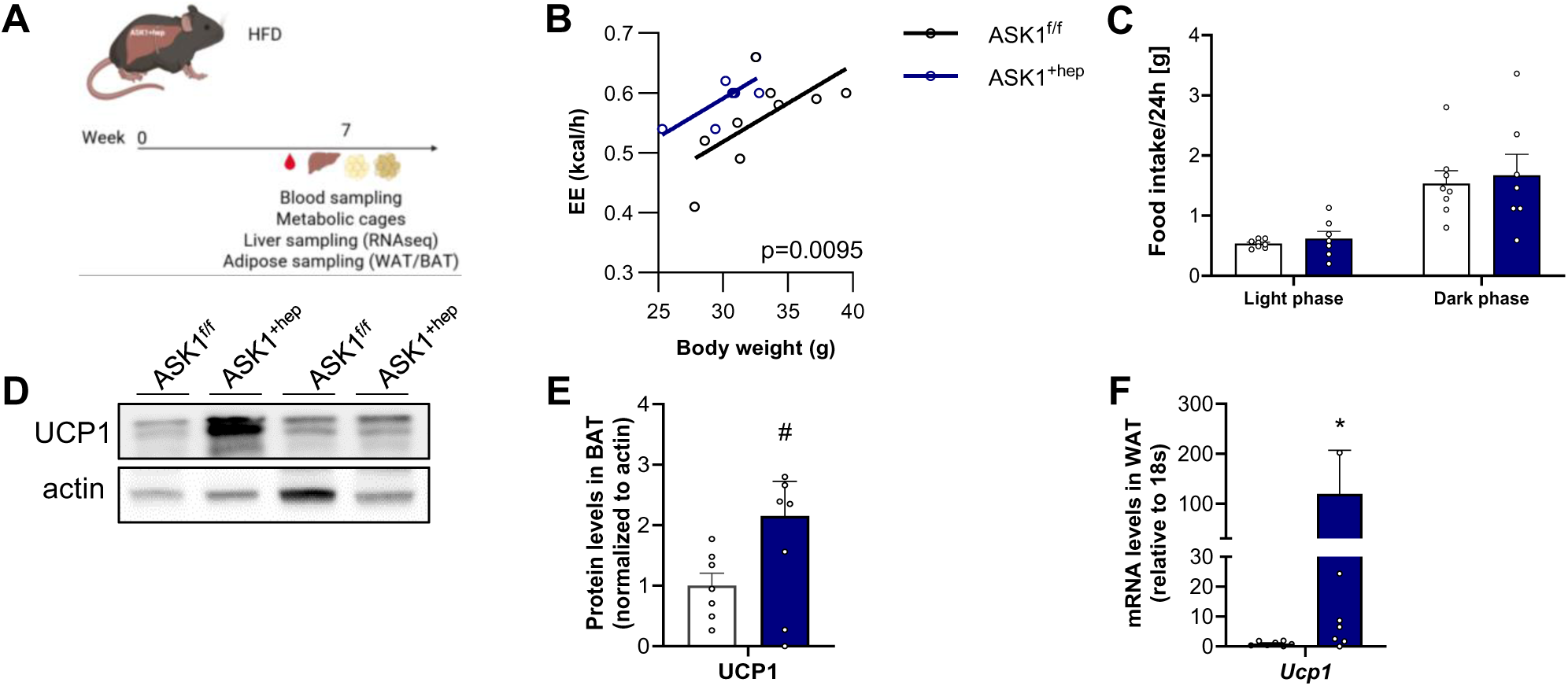
Energy expenditure is increased in HFD-fed ASK1^+hep^ mice. **A**. Schematic of experimental procedures performed in mice fed a HFD for 7 weeks. **B.** Linear regression analysis of energy expenditure in ASK1^+hep^ and ASK1^f/f^ mice as a function of body mass (n=… mice). **C.** Food intake in ASK1^+hep^ (blue bars) and ASK1^f/f^ (white bar) mice on HFD for 7 weeks during light phase and dark phase (n=6-8 mice). **D.** Representative Western blot of UCP1 and actin in BAT of ASK1^f/f^ and ASK1^+hep^ mice. **E.** Protein quantification of UCP1 (normalized to actin) in BAT of ASK1^+hep^ (blue bars) and ASK1^f/f^ (white bars) mice (n=7-8). **F.** mRNA expression of *Ucp1* in inguinal white adipose tissue (ingWAT) of ASK1^+hep^ (blue bars) and ASK1^f/f^ (white bars) mice (n=7-8). Data are shown as mean ± SEM. ANCOVA for B. ^#^p<0.1, *p<0.05 (Mann-Whitney test for panel E and F).

In line, levels of the master regulator of non-shivering thermogenesis - UCP1 - were increased in BAT and ingWAT in ASK1^+hep^ mice after 7 weeks of HFD (Figs. 2 D-F).

### FGF21 plasma concentration is elevated in HFD-fed ASK1^+hep^ mice

Next, we performed bulk RNA-sequencing of the liver to identify potential hepatokines that mediate increased energy expenditure in 7-week HFD ASK1^+hep^ mice (Fig. 2A). Of note, liver, total white adipose tissue and body weights were not significantly different between the two genotypes after 7 weeks of HFD (Fig. 1A and Suppl. Figs. 2 A-C). Differential gene expression analysis revealed *Fgf21* to be the most highly upregulated gene in livers of ASK1^+hep^ mice (Fig. 3A). In line, circulating FGF21 levels were significantly increased in 7-week HFD-fed ASK1^+hep^ mice (Fig. 3B). In agreement, plasma FGF21 levels were significantly higher in 20-week HFD-fed ASK1^+hep^ mice (Fig. 3C), and the latter correlated negatively with body weights (Fig. 3D). In contrast, chow-fed ASK1^+hep^ mice did not display differences in circulating FGF21 levels compared to littermate controls (Suppl. Fig. 2D). Taken together, our data support the notion that elevated FGF21 levels in HFD-fed ASK1^+hep^ mice critically contribute to the reduced body weight gain in these mice.

**Figure 3.**
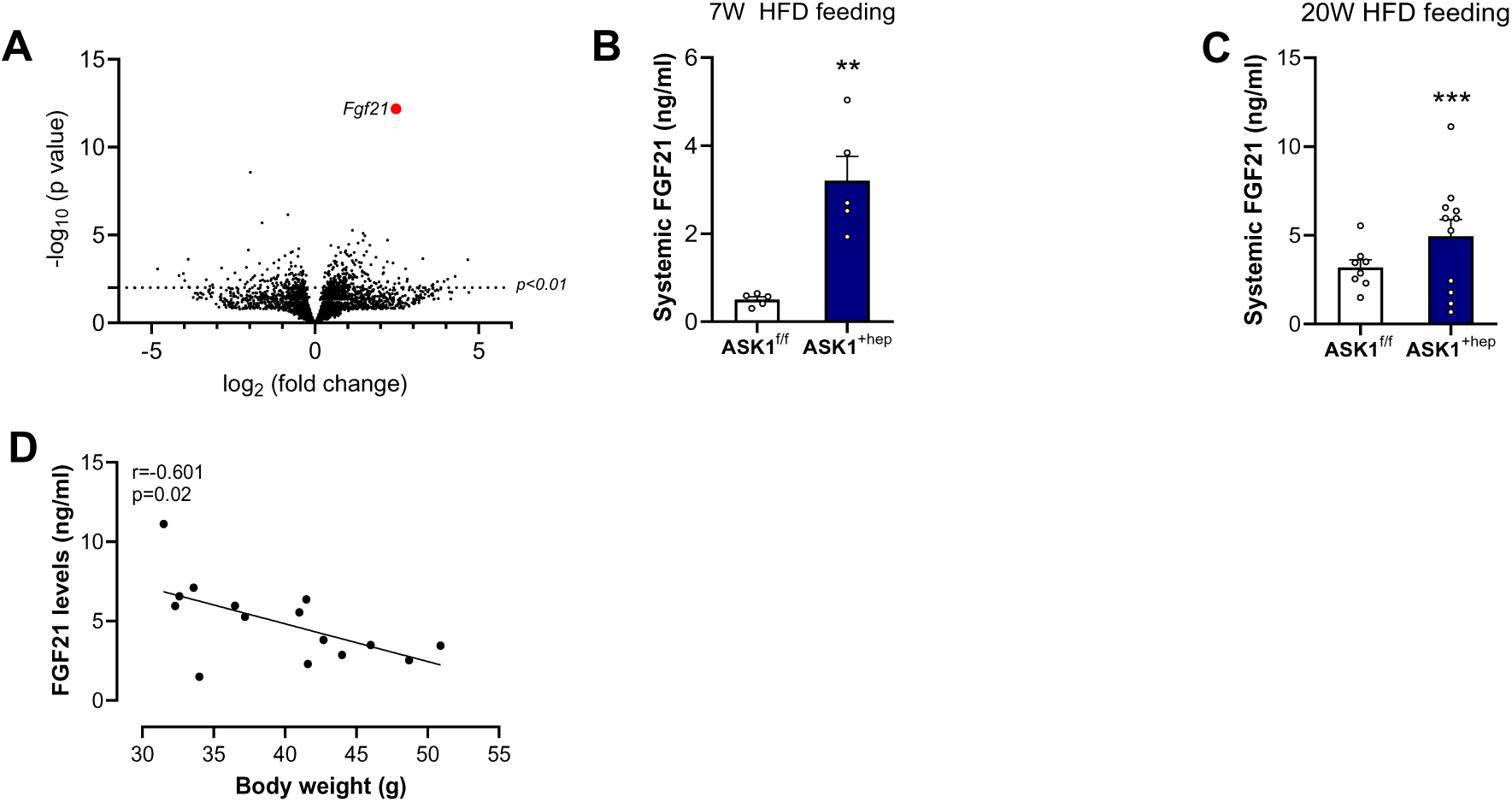
FGF21 plasma concentration is elevated in HFD-fed ASK1^+hep^ mice. **A.** Volcano plot of hepatic RNA sequencing data of ASK1^+hep^ and ASK1^f/f^ mice after 7 weeks on HFD (n=7-8 mice). **B**. Circulating FGF21 plasma levels in HFD-fed ASK1^+hep^ (blue bars) and ASK1^f/f^ (white bars) mice after 7 weeks on HFD (n=5 mice per group). **C**. Circulating FGF21 plasma levels in HFD-fed ASK1^+hep^ (blue bars) and ASK1^f/f^ (black bars) mice after 20 weeks on HFD (n=8-10). **D.** Correlation between body weight and circulating FGF21 plasma levels in 20W HFD-fed ASK1+hep and ASK1f/f mice (n=15). Data are shown as mean ± SEM. **p<0.01, ***p<0.001 (Student’s t test for panels B and C, Pearson’s correlation coefficient r for panel D).

### Increased FGF21 expression in HFD-fed ASK1^+hep^ mice is mediated by ATF4

ASK1 is a serine/threonine protein kinase and a member of the MAPK kinase kinase (MAP3Ks) family. Upon activation, ASK1 undergoes homodimerization and autophosphorylation and subsequently induces phosphorylation and activation of downstream kinases such as c-Jun N-terminal kinase (JNK) and p38 MAPK (23–25). To investigate whether these kinases contribute to increased *Fgf21* expression in livers of HFD-fed ASK1^+hep^ mice, experiments in primary hepatocytes treated with JNK or p38 inhibitors were performed (Fig. 4A). p38 inhibitor treatment of ASK1^+hep^-derived primary hepatocytes reduced *Fgf21* transcript levels to a significantly higher extent than in ASK1^f/f^ derived primary hepatocytes (Fig. 4B and Suppl Fig 3A). In contrast, inhibiting JNK in ASK1^+hep^ hepatocytes did not reduce *Fgf21* expression levels (Suppl Fig 3B). This data indicates that p38 rather than JNK mediates the effect of ASK1 overexpression on *Fgf21* transcription in hepatocytes. To affect transcription, kinases such as p38 (24, 25) activate transcription factors, which eventually change gene expression by binding to the promoter of respective genes. To identify potential transcription factors that mediate the effect of the ASK1-p38 axis on the expression of *Fg21*, we performed transcription factor enrichment analysis using liver RNA-sequencing data (Fig. 3A), using ChIP-X Enrichment Analysis 3 (ChEA3). ChEA3 is a computational tool that ranks transcription factors associated with user-submitted gene sets based on evidence from multiple experimental and computational datasets (26). As shown in Fig. 4C, transcriptional activity of the activating transcription factor 4 (ATF4) was elevated among other factors in livers of HFD-fed ASK1^+hep^ mice. Importantly, ATF4 can be activated by p38 (27). Moreover, recent evidence indicates that the stress-induced increase in *Fgf21* expression is mediated by ATF4 (28, 29). To test the hypothesis whether ATF4 is mediating ASK1-p38-induced *Fgf21*-expression, ATF4 was depleted in primary hepatocytes isolated from HFD-fed ASK1^f/f^ and ASK1^+hep^ mice. Hepatocytes were treated with scrambled siRNA or siRNA targeting ATF4. As expected, siRNA-mediated knockdown of ATF4 significantly downregulated *Atf4* expression in both genotypes (Suppl. Fig. 3C). Moreover, increased *Ask1* expression in ASK1^+hep^ mice (Suppl. Fig. 3D) was paralleled by significantly increased *Fgf21* transcription in hepatocytes treated with scrambled siRNA (Fig. 4D). Importantly, ATF4 depletion significantly downregulated *Fgf21* in hepatocytes derived from ASK1^+hep^ but not from ASK1^f/f^ mice (Fig. 4D), indicating that ATF4 plays a crucial role in the upregulation of *Fgf21* expression in ASK1 overexpressing hepatocytes.

**Figure 4.**
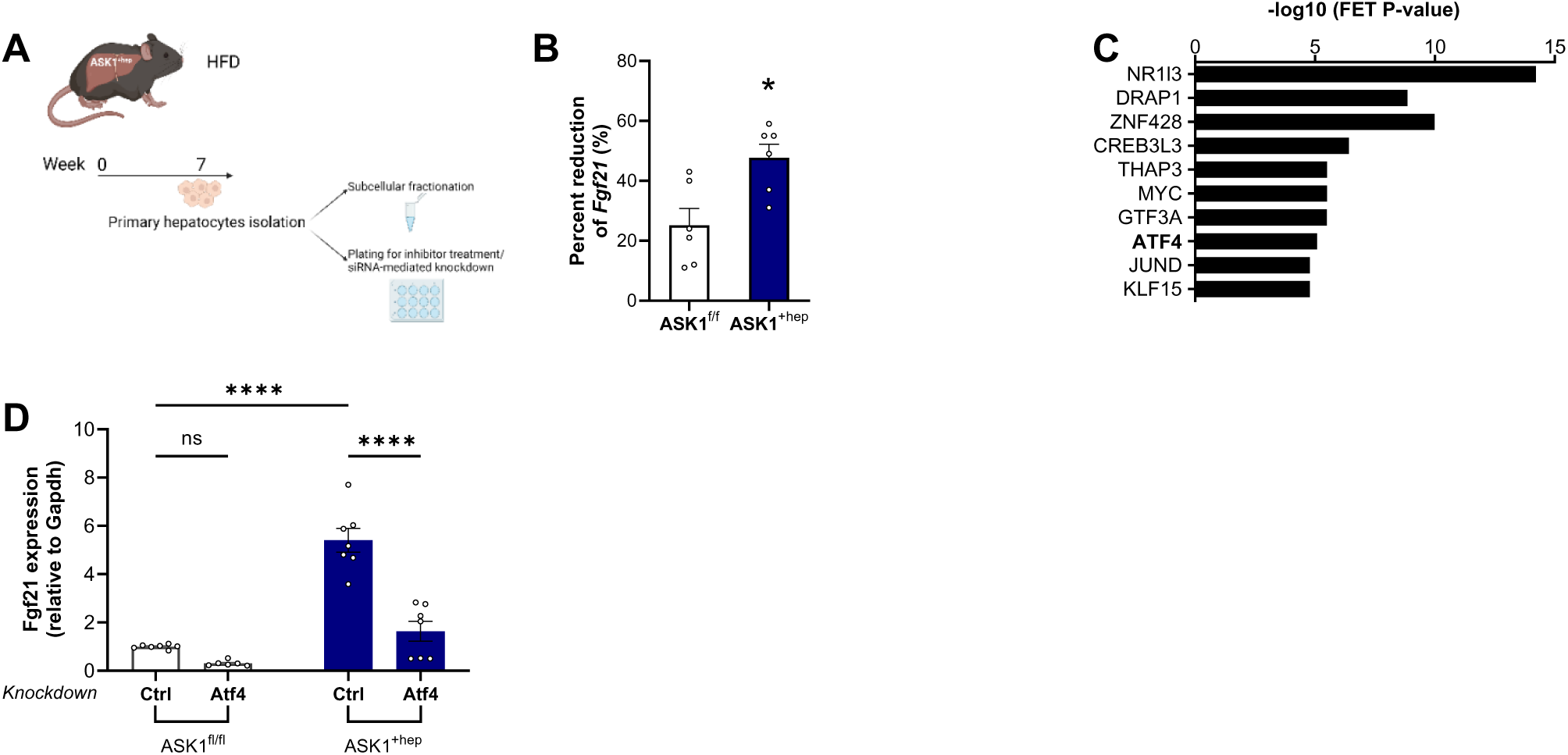
Increased FGF21 expression in HFD-fed ASK1^+hep^ mice is mediated by ATF4. **A**. Schematic of experimental design after isolation of primary hepatocytes from HFD-fed ASK1^fl/fl^ and ASK1^+hep^ mice **B.** Percent reduction of *Fgf21* gene expression after p38-inhibitor treatment of primary hepatocytes isolated from 7 weeks HFD-fed ASK1^+hep^ (blue bar) and ASK1^f/f^ (white bar) mice (n=6 mice per group). **C**. Top 10 enriched transcription factors in 7 weeks HFD-fed ASK1^+hep^ vs ASK1^f/f^ livers. Data are presented as -log10 of the FET p-value. **D.** *Fgf21* gene expression after siRNA-mediated knockdown of ATF4 in primary hepatocytes isolated from 7 weeks HFD-fed ASK1^+hep^ and ASK1^f/f^ mice (n=5-6). Data are shown as mean ± SEM. *p<0.05, ****p<0.0001 (Two-way ANOVA for panels B and E, Student’s t test for panels C and F).

## 3. Discussion

Obesity and metabolic dysfunction is associated with activation of stress-activated protein kinases such as ASK1 (30). In line, we previously identified a protective role of hepatic ASK1 in the development of obesity-induced MAFLD (22). In particular, we found that liver-specific ASK1 knockout aggravated HFD-induced hepatic steatosis, whereas liver-specific ASK1-overexpressing mice (ASK1^+hep^) revealed reduced hepatic MAFLD. Herein, we report that liver-specific ASK1 overexpression prevented HFD-induced obesity and associated deterioration of glucose metabolism. This lean and healthy metabolic phenotype of HFD-fed ASK1^+hep^ mice is presumably mediated by FGF21, given that HFD-fed ASK1^+hep^ mice displayed significantly higher hepatic *Fgf21* expression as well as elevated systemic FGF21 levels. Moreover, resistance to diet-induced obesity in ASK1^+hep^ mice conceivably resulted from increased energy expenditure, likely driven, at least in part, by elevated thermogenesis in BAT and WAT.

Of note, ATF4 is known to be a transcriptional regulator of *Fgf21* gene transcription in response to nutritional stimuli (31). Mechanistically, we speculate that the transcription factor ATF4 may be induced by HFD-induced ASK1-p38 signaling in hepatocytes, and in turn stimulates hepatic *Fgf21* gene expression by binding to the respective promoter region. In support of our hypothesis, p38 inhibition reduced *Fgf21* expression to a higher extent in ASK1 overexpressing compared to control hepatocytes. Moreover, silencing ATF4 in cultured primary hepatocytes significantly reduced *Fgf21* transcript levels in cells derived from ASK1^+hep^ but not ASK1^fl/fl^ mice

It remains to be understood how exactly liver-expressed ASK1 leads to increased production and/or secretion of FGF21. We propose that ATF4 mediates ASK1-induced *Fgf21* expression. ATF4 is an adaptive response regulator of metabolic homeostasis and plays a key role in both the adaptation to cellular stress and the activation of apoptosis (32–34). Once activated, ATF4 translocate to the nucleus and binds to the promoter regions of different genes to induce changes in gene expression (34). Based on the *in silico* analysis of the bulk RNA sequencing data, ATF4 appeared in the top 10 enriched transcription factors that are predicted to play a role based on the differential expressed genes. Indeed, ATF4 is known as a transcriptional regulator of *Fgf21* gene expression in response to nutritional cues such as high fat diets (31). In addition, an interaction between the MAP kinase p38, a downstream kinase of ASK1, and ATF4 has previously described (27, 35). Hence, we hypothesized that ASK1 overexpression activated the p38-ATF4 axis, which in turn induces *Fgf21* gene expression.

In nutritional or genetic mouse models of obesity, administration of bio-engineered FGF21 enhances energy expenditure, while it confers resistance to HFD-induced obesity and improves glucose tolerance and whole-body insulin sensitivity (7, 8, 36). Hence, the favorable metabolic phenotype in HFD-fed ASK1^+hep^ mice likely derives from increased energy expenditure, driven by elevated circulating FGF21 concentrations. Mechanistically, it was proposed that FGF21 activates thermogenesis by activating brown adipose tissue and inducing the browning of white adipose tissue. In line, HFD-fed ASK1^+hep^ mice showed increased UCP1 levels in brown and white adipose tissue. Yet, we cannot exclude FGF21-mediated, UCP1-independent, effects affecting energy balance in ASK1^+hep^ mice as has been previously suggested (37–39). Moreover, other energy-consuming processes, such as futile substrate cycling, in adipose tissue might additionally contribute to the observed increase in energy expenditure HFD-fed ASK1^+hep^ mice.

Given the role of ASK1 as a signaling node for stress signals and its central role in inflammatory processes (30), tissue-specific overexpression of ASK1 should intuitively lead to harmful effects in the affected tissue. Accordingly, Xiang et al. showed that liver-specific ASK1 transgenic mice exhibited aggravated obesity and insulin resistance compared with controls in response to HFD administration (40). Yet, drug candidates such as the ASK1 inhibitor Selonsertib have so far failed to successfully treat liver fibrosis in MASH patients, thus challenging this treatment concept (Harrison et al. 2020). In addition, we didn’t observe an upregulation of inflammatory markers or stress-induced harm in livers of HFD-fed ASK1^+hep^ mice (22).The recent identification of liver-expressed ASK1 as a protective factor in the development of obesity-induced MAFLD by Challa et al. further questions the ascribed detrimental role of ASK1 in liver-associated metabolic disease (22). Thus, it is challenging to explain the discrepancy between the phenotype we observe in our study and the phenotype reported in the article by Xiang et al. In both set-ups, liver-specific ASK1 overexpression was mediated by constitutive overexpression of ASK1 (as opposed to inducible overexpression), mice were subjected to long-term HFD exposure (20 vs. 24 weeks) and housing conditions (most importantly housing temperatures) were comparable. Moreover, liver-specific ASK1 overexpression yielded a 2-2.5-fold overexpression in both mouse models. However, there are at least two conceivable differences that might, at least partly, explain the diametrically opposed phenotypes. First, the composition of the high-fat diet used for the experiments was different. While the fat fraction in the diet of the present study was based on coconut oil, the HFD used by Xiang et al. was lard-based. We speculate that the different fatty acid composition of the diets may account for some of the different outcomes that we find. The individual fatty acids derived from different fat sources may mediate very different, direct effects on affected tissues or on processes such as inflammation (41, 42). Secondly, the discrepancies could arise from differences in the genetic approach to yield ASK1 overexpression. Xiang et al. generated a mouse model with a constitutively active ASK1 kinase in hepatocytes (40). In our study, we employed a mouse model in which the stop-cassette in front of the inserted ASK1 cDNA was excised by *Cre* recombinase. However, hepatic ASK1 was not constitutively active in our mouse model. We speculate that the activity of liver-expressed ASK1 in our mice is subjected to environmental cues such as light phase or feeding and thus enhanced ASK1 activity fluctuated in response to circadian rhythms. The fact that chow-fed ASK1^+hep^ mice did not have a different metabolic phenotype compared to *Cre*-negative littermate mice supports the idea that ASK1 activity is indeed triggered by environmental cues (in our case HFD) and only in such a situation exerts its effects. Hence, it might be the difference between constitutive ASK1 activity in the study of Xiang et al. and the dynamic activity of ASK1 in our study that explains the discrepancies in the observed phenotype in HFD-fed liver-specific ASK1 overexpressing mice.

Of note, we found no significant effect of ASK1 overexpression on circulating FGF21 concentrations, body weight gain and glucose metabolism in chow-fed mice. This finding suggests that hepatocyte-specific ASK1 overexpression alone has no overt effect on hepatic *Fgf21* expression and, hence, energy and glucose homeostasis. Instead, a metabolic stressor, such as a fat-enriched diet, may be required to activate ASK1 signaling and, thus, induce *Fgf21* gene expression in ASK1^+hep^ mice. Indeed, it has been reported that in absence of strong stimuli, for example when exposed to a regular rodent diet, ASK1 activation is blocked by binding of thioredoxin to the N-terminus of ASK1 (43). The observation that hepatic ASK1 overexpression only elevates FGF21 concentrations upon HFD exposure may highlight the evolutionary importance of FGF21 as a hormone protecting against nutritional stressors. To our knowledge, we are the first group reporting that hepatic *Fgf21* expression is induced by the stress kinase ASK1 upon HFD-feeding, thereby lowering body weight and preserving metabolic health. However, the notion that ASK1 can induce FGF21 expression is not completely novel. In 2021, Ogawa et al. reported that ASK1-p38 signaling induces FGF21 to promote mechanical cell competition and motility in a mammalian cell line called Madin-Darby canine kidney cells (44). In this context, the ASK1-FGF21 axis regulates cell-cell communication and tissue maintenance, and it is likely to assume that the involvement of FGF21 in cell motility is due to the fact that FGF21 is also a growth factor besides being a metabolically active hormone. In contrast, we uncovered a mechanistic link between liver-expressed ASK1 and metabolically active FGF21, which regulates energy expenditure and, thus, body weight in mice.

FGF21 has gained a lot of scientific interest in order to decipher its mechanism of action and to elucidate ways to leverage its circulating concentrations as to use it as a therapeutic agent. Indeed, there are on-going attempts to increase circulating FGF21 levels to alleviate obesity and metabolic diseases, mostly by injection of recombinant FGF21 or FGF21 analogues. While this strategy works reasonably well in mice, it has had limited success in obese primates and human subjects, mainly due to adverse side effects and a short half-life times (45, 46). Herein, we have generated a mouse model in which overexpression of the stress kinase ASK1 in the liver leads to endogenously elevated FGF21 concentrations. Rather than acutely elevating circulating FGF21 via injection of a recombinant protein, continuously increasing circulating FGF21 by elevating endogenous production may be a promising approach to circumvent drawbacks of exogenous FGF21. We hypothesize that this concept would allow to avoid two main limitations of recombinant FGF21 or FGF21 analogues: adverse side effects such as elevation of heart rate and blood pressure as well as gastrointestinal disorders (47, 48) as well as problems provoked by the short half-life time of injected FGF21 (45).

In conclusion, we have uncovered a yet undescribed regulatory axis between hepatic ASK1 and FGF21 which positively affects the regulation of energy and glucose metabolism. Given the prominent role of FGF21 in whole-body metabolism and body weight control, the identification of ASK1 as a potential cell-intrinsic *Fgf21* enhancer may contribute to the development of new drugs counteracting obesity and associated diseases.

---

**Supplementary Figure 1.**
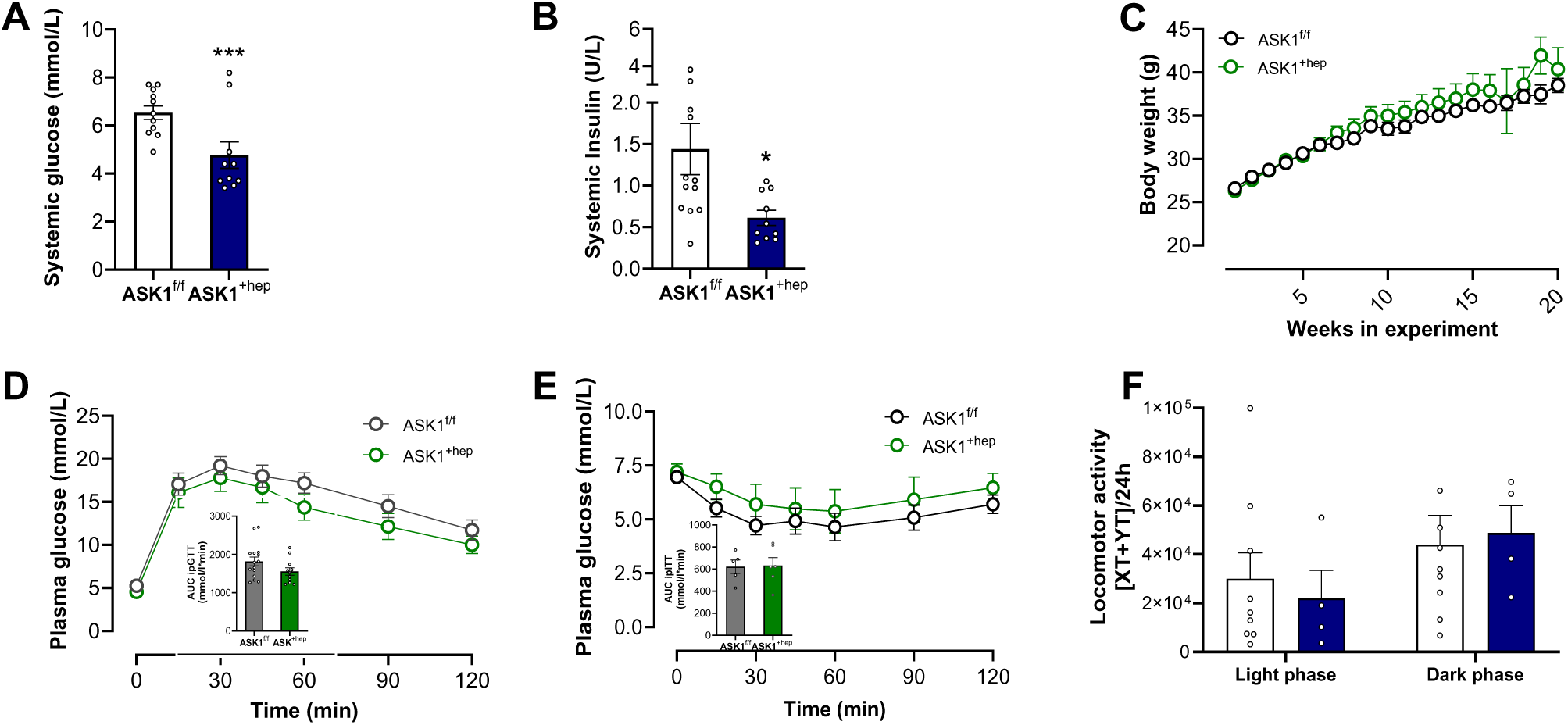
**A.** Plasma insulin levels after 5 h fasting in ASK1^+hep^ and ASK1^f/f^ mice on HFD for 20 weeks (n=10-11 mice). **B.** Plasma glucose levels after 5h fasting in ASK1^+hep^ and ASK1^f/f^ mice on a HFD for 20 weeks (n=10-11 mice). **C.** Body weight development over time of chow-fed ASK1^f/f^ and ASK1^+hep^ mice (n=12-13). **D.** ipGTT (including area under the curve (AUC)) in ASK1^+hep^ and ASK1^f/f^ mice fed a chow diet (CD) for 20 weeks (n=6-10 mice). **E.** ipITT (including area under the curve (AUC)) in ASK1^+hep^ and ASK1^f/f^ mice fed a CD for 20 weeks (n=5-7 mice). **F.** Locomotor activity of ASK1^+hep^ (blue bar) and ASK1^f/f^ (white bar) mice on HFD for 7 weeks during light phase and dark phase (n=6-8 mice). Data are shown as mean ± SEM. **p<0.01, ***p<0.001 (Student’s *t* test for panel A).

**Supplementary Figure 2.**
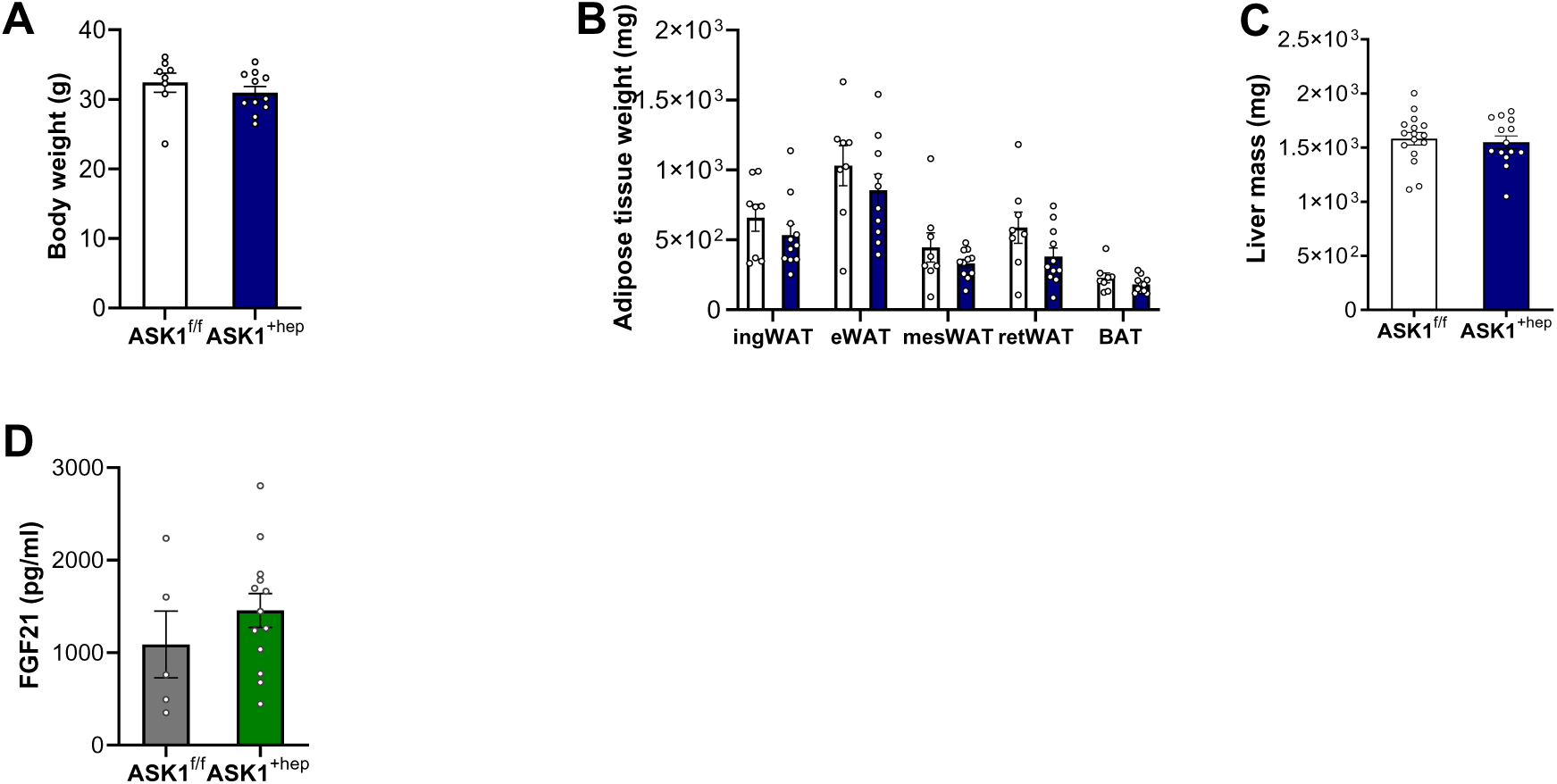
**A**. Body weights of ASK1^f/f^ and ASK1^+hep^ mice fed a HFD for 7 weeks (n=8-11 mice). **B.** Weight of different fat pads of ASK1^f/f^ and ASK1^+hep^ mice fed a HFD for 7 weeks (n=8-11 mice); inguinal (ingWAT), epididymal (eWAT), mesenteric (mesWAT), retroperitoneal (retWAT) and brown adipose tissue (BAT). **C.** Liver weight of ASK1^f/f^ and ASK1^+hep^ mice fed a HFD for 7 weeks (n=14-16 mice). **D.** FGF21 plasma concentration in 26 weeks old chow-fed ASK1^+hep^ and ASK1^f/f^ mice (n=5-13). Data are shown as mean ± SEM.

**Supplementary Figure 3.**
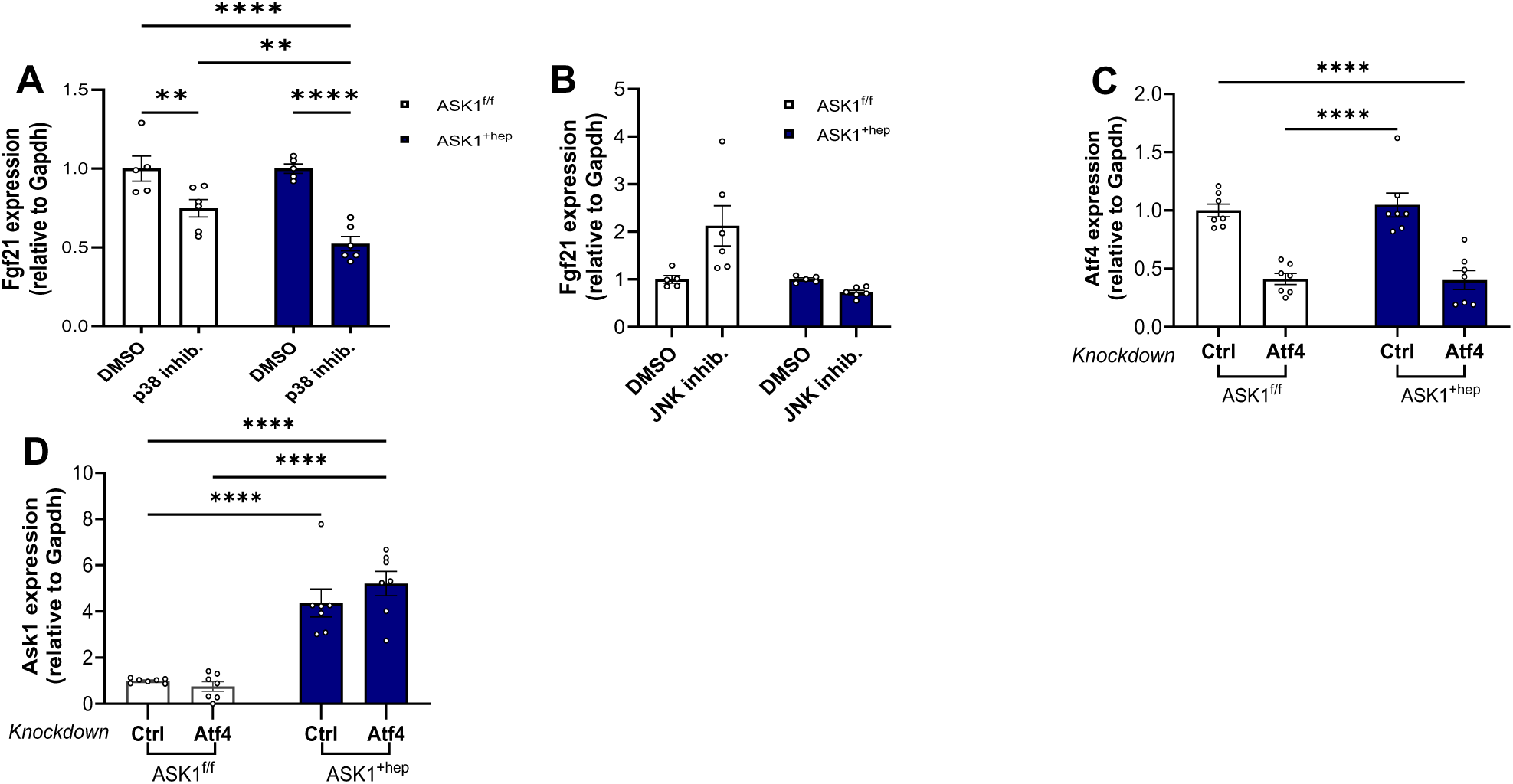
**A.** *Fgf21* gene expression after treating primary hepatocytes isolated from 7 weeks HFD-fed ASK1^+hep^ and ASK1^f/f^ mice with DMSO or p38 inhibitor for 24h (n=6 mice). **B**. Percent reduction of *Fgf21* gene expression after JNK-inhibitor treatment of primary hepatocytes isolated from 7 weeks HFD-fed ASK1^+hep^ and ASK1^f/f^ mice (n=6) **C.** *Atf4* gene expression after siRNA-mediated knockdown of ATF4 in primary hepatocytes isolated from 7 weeks HFD-fed ASK1^+hep^ and ASK1^f/f^ mice (n=6). **D.** *Ask1* gene expression after siRNA-mediated knockdown of ATF4 in primary hepatocytes isolated from 7 weeks HFD-fed ASK1^+hep^ and ASK1^f/f^ mice (n=6). Data are shown as mean ± SEM. **p<0.01, ***p<0.001, **** p<0.0001 (Two-way ANOVA for panel A-D).

## Material and Methods

### Animal husbandry and experiments

Liver-specific ASK1-overexpressing (ASK1^+hep^) mice on a C57BL/6 background were generated as previously described (Challa et al. 2019). As controls, floxed littermates (ASK1^f/f^) were used. All animal experiments were conformed to the Swiss animal protection laws and were approved by the Cantonal Veterinary Office in Zurich, Switzerland (Licenses ZH193/2017 and ZH103/2021). Male mice, aged six to 26 weeks, were used for studies. All experiments were performed with mice kept in a 12h: 12h light: dark cycle (light phase starting at 7 a.m.) in a pathogen-free animal facility. Cages were enriched with crinklets and gentle tunnel handling was used. Animals were monitored weekly (habitus, fur, activity, locomotion, eye symptoms, body weight). Human endpoints were defined as signs of pain (humpback or closed eyes) or body weight loss >10%. Group allocation regarding diet (chow vs. HFD) was based on the initial body weight (similar average body weight in both groups). Group size was determined based on previous experiments performed in our laboratory. Experimenters were not blinded to group allocations. Mice were housed 2-5 mice per cage, in individually ventilated cages. The ambient temperature in the animal facility was kept constant at 22°C and the animals were fed standard rodent (chow) diet or 59 % HFD (E15772-347, ssniff-Spezialdiäten GmbH, Soest, Germany), with ad libitum access to food and water. HFD experiments started when mice were 6 weeks old, and maintained for either 7 or 20 weeks. Tissue was collected in animals fasted for 5 hours (from 8 am to 1 pm).

### Isolation of primary hepatocytes

Mice were euthanized by CO_2_ asphyxiation and the abdominal cavity opened to expose the vena cava inferior. The latter was cannulated with a catheter and the liver perfused with 1x HBSS supplemented with 0.05 mM EDTA for 4 minutes, followed by a digestion using DMEM (Glucose 1 g/L), containing 1% P/S, 15 mM HEPES, and 32 µg/L Liberase^TM^ for 4 minutes. The liver was removed from the mouse and cells released from the liver into ice-cold DMEM (Glucose 1 g/L) supplemented with 10 % FBS and 1 % P/S (full medium) until only connective tissue was left. The cells were filtered through a 100 µM filter and washed 3 times in full medium with intermittent centrifugations at 50 g, 4°C and 2 min. Primary hepatocytes were isolated from the remaining cells by centrifugation with 90% Percoll and viable cells plated in plating medium at a density of 2*10^5^ cells/ml as described (Jung, Zhao, and Svensson 2020).

### siRNA-mediated knockdown

After plating, primary hepatocytes were allowed to attach and recover for 3 hours. Before transfection, plating medium was replaced by maintenance medium (Williams E medium, prepared as described in (Jung, Zhao, and Svensson 2020). For RNA interference, Opti-Mem and 200 nM small interfering RNA (siRNA) targeting ATF4 (siATF4; ON-TARGETplus Mouse Atf4 siRNA, L-042737-01-0005, Horizon Discovery Biosciences Limited, Cambridge, UK) or control siRNA (siCtrl; ON-TARGETplus Non-targeting Pool, D-001810-10-05; Horizon Discovery Biosciences Limited) were mixed and primary hepatocytes were transfected using Lipofectamine 2000 with antibiotic-free maintenance medium. After 24 h, medium was aspirated, cells washed with ice-cold PBS and frozen −80°C until further processing.

### RNA isolation and quantitative RT-PCR

Total RNA was extracted with the NucleoSpin® RNA isolation kit (Macherey-Nagel, Düren, Germany). RNA concentration was determined using NanoDrop® spectrophotometer (ThermoFisher Scientific). Equal amounts of RNA were reverse-transcribed with the GoScript™ Reverse Transcription System (Promega) and cDNA amplified by TaqMan real-time PCR using the following probes/primers: Ask1, Mm00434883_m1; Atf4, Mm00515324_m1; Fgf21, Mm00840165_m1; Gapdh, Mm99999915_g1. (Applied Biosystems, Rotkreuz, Switzerland). Relative gene expression values were obtained after normalization to Gapdh using the 2^−ΔΔCt^ method. (Pfaffl 2001).

### RNA sequencing and transcription factor enrichment analysis

RNA sequencing was conducted by Novogene (Cambridge, UK). mRNA was purified using poly-T oligo-attached magnetic beads. cDNA synthesis was performed using random hexamer primers, followed by library preparation with specific protocols for directional and non-directional libraries, including end repair, A-tailing, adapter ligation, size selection, and amplification. Libraries were quantified using Qubit, real-time PCR, and bioanalyzer before pooling and sequencing on Illumina platforms. Raw fastq data was processed to obtain clean reads by removing adapters, low-quality reads, and calculating Q20, Q30, and GC content. Clean reads were mapped to the reference genome using Hisat2, with read counts quantified by FeatureCounts and gene expression estimated as FPKM. Differential expression analysis was conducted with DESeq2 for biological replicates and edgeR for non-replicates, applying Benjamini-Hochberg corrections. Genes with adjusted p-values ≤ 0.05 were deemed differentially expressed. Transcription factor analysis was performed using ChEA3 to identify enriched transcription factors (Keenan et al. 2019).

### Protein quantification and Western blot analysis

Tissue or cells were lysed in RIPA buffer 150 mM NaCl, 50 mM Tris-HCl (pH 7.5), 1 mM EGTA, 1% NP-40, 0.25% sodium deoxycholate, 1 mM sodium pyrophosphate, 1 mM sodium vanadate, 1 mM NaF, 10 mM sodium β-glycerophosphate, 0.2 mM PMSF and a 1:1000 dilution of protease inhibitor cocktail (Sigma-Aldrich, Saint-Louis, MI, USA). The protein concentration in each sample was quantified using the BCA protein assay kit (Pierce, Rockford, IL, USA). Equal amounts of proteins (50 μg) were separated on a 12% polyacrylamide gel using SDS-PAGE and electro-transferred to a nitrocellulose membrane (0.2 μm, BioRad, Reinach, Switzerland). Ponceau S staining was used to confirm equal protein loading on membranes. The membranes were then blocked in 5% non-fat dry milk dissolved in Tris-buffered saline (50 mM Tris-HCl, 150 mM NaCl) containing 0.1% Tween (TBS-T) and subsequently incubated with a primary antibody overnight at 4 °C. The following day, membranes were incubated with corresponding secondary HRP–conjugated antibodies. Antibody-antigen complexes were detected by using ChemiDoc Imaging System (Bio-Rad Laboratories). The following primary antibodies were used: UCP1, PA1-24894 (ThermoFisher Scientific, Waltham, MA, USA; diluted 1:1000); Actin, MAB1501 (Millipore, Darmstadt, Germany; diluted 1:5000). Protein levels were quantified using the Image Lab software (BioRad, version 5.2.1) and normalized to Actin.

### Circulating FGF21 and insulin levels

Plasma FGF21 and insulin levels were measured using the kits *Mouse and rat FGF21 ELISA* (Biovendor) and the Insulin Mouse Ultra Sensitive ELISA (Chrystal Chem), respectively.

### Indirect calorimetry

Indirect calorimetry measurements were performed with the opencircuit Phenomaster system (TSE Systems) according to the manufactureŕs guidelines and protocols. Animals were single-caged and acclimated to the metabolic cage for 24 hours before starting the measurements for 72 hours. Food intake, locomotor activity as well as O_2_ consumption and CO_2_ production were recorded throughout the whole measurement. Locomotor activity was measured using a 2-dimensional infrared light-beam. O_2_ and CO_2_ levels were recorded 3 times per hour. Energy expenditure was calculated according to the manufactureŕs guidelines. Individual energy expenditure and food intake data points shown in graphs correspond to the average value of 72h for each mouse.

### Insulin and glucose tolerance test

For intraperitoneal insulin tolerance tests (ipITT), animals were fasted for 3h (from 9 am to 12 pm) and for intraperitoneal glucose tolerance tests (ipGTT) overnight (from 5 pm to 8 am). Blood glucose concentration was measured before injection of human recombinant insulin (1.0 U/kg body weight) or glucose (2 g/kg body weight) and 15, 30, 45, 60, 90 and 120 min thereafter. Blood glucose concentrations were measured with a blood glucometer (Accu-Chek Aviva; Roche Diagnostics) in blood collected from tail vein incisions.

### Data analysis

All results are expressed as mean ± standard error of the mean (SEM). Statistical analysis was performed using two-tailed, unpaired Student’s *t*-test or two-way ANOVA with Bonferroni multiple comparisons (GraphPad Prism Software, San Diego, CA, USA; Version 8.0.0), assuming normal distribution. If the data were not normally distributed, Mann-Whitney test was performed. Data were not included if misinjection occurred during ITT/GTT. Outliers defined by the ROUT test were excluded from statistical analysis. p values < 0.05 were considered to be statistically significant. GraphPad was used for statistical analysis and to produce graphs; BioRender was used to create figures.

## Literature

1. Tremmel M, Gerdtham U-G, Nilsson PM, Saha S. Economic Burden of Obesity: A Systematic Literature Review. Int J Environ Res Public Health. 2017;14(4):435.

2. Washington TB, Johnson VR, Kendrick K, Ibrahim AA, Tu L, Sun K, et al. Disparities in Access and Quality of Obesity Care. Gastroenterol Clin North Am. 2023;52(2):429–41.

3. Zhang X, Ha S, Lau HC, Yu J. Excess body weight: Novel insights into its roles in obesity comorbidities. Semin Cancer Biol. 2023;92:16–27.

4. Saklayen MG. The Global Epidemic of the Metabolic Syndrome. Curr Hypertens Rep. 2018;20(2):12.

5. Virani SS, Alonso A, Aparicio HJ, Benjamin EJ, Bittencourt MS, Callaway CW, et al. Heart Disease and Stroke Statistics-2021 Update: A Report From the American Heart Association. Circulation. 2021;143(8):e254–e743.

6. Markan KR, Naber MC, Ameka MK, Anderegg MD, Mangelsdorf DJ, Kliewer SA, et al. Circulating FGF21 Is Liver Derived and Enhances Glucose Uptake During Refeeding and Overfeeding. Diabetes. 2014;63(12):4057–63.

7. Kharitonenkov A, Shiyanova TL, Koester A, Ford AM, Micanovic R, Galbreath EJ, et al. FGF-21 as a novel metabolic regulator. J Clin Invest. 2005;115(6):1627–35.

8. Coskun T, Bina HA, Schneider MA, Dunbar JD, Hu CC, Chen Y, et al. Fibroblast growth factor 21 corrects obesity in mice. Endocrinology. 2008;149(12):6018–27.

9. Xu J, Lloyd DJ, Hale C, Stanislaus S, Chen M, Sivits G, et al. Fibroblast growth factor 21 reverses hepatic steatosis, increases energy expenditure, and improves insulin sensitivity in diet-induced obese mice. Diabetes. 2009;58(1):250–9.

10. Fisher FM, Maratos-Flier E. Understanding the Physiology of FGF21. Annu Rev Physiol. 2016;78:223–41.

11. Engin A. Reappraisal of Adipose Tissue Inflammation in Obesity. Adv Exp Med Biol. 2024;1460:297–327.

12. Hondares E, Iglesias R, Giralt A, Gonzalez FJ, Giralt M, Mampel T, et al. Thermogenic activation induces FGF21 expression and release in brown adipose tissue. J Biol Chem. 2011;286(15):12983–90.

13. Chartoumpekis DV, Habeos IG, Ziros PG, Psyrogiannis AI, Kyriazopoulou VE, Papavassiliou AG. Brown adipose tissue responds to cold and adrenergic stimulation by induction of FGF21. Mol Med. 2011;17(7-8):736–40.

14. Fisher FM, Kleiner S, Douris N, Fox EC, Mepani RJ, Verdeguer F, et al. FGF21 regulates PGC-1α and browning of white adipose tissues in adaptive thermogenesis. Genes Dev. 2012;26(3):271–81.

15. Cannon B, Nedergaard J. Brown adipose tissue: function and physiological significance. Physiol Rev. 2004;84(1):277–359.

16. Talukdar S, Zhou Y, Li D, Rossulek M, Dong J, Somayaji V, et al. A long-acting FGF21 molecule, PF-05231023, decreases body weight and improves lipid profile in non-human primates and type 2 diabetic subjects. Cell metabolism. 2016;23(3):427–40.

17. Tillman EJ, Rolph T. FGF21: An Emerging Therapeutic Target for Non-Alcoholic Steatohepatitis and Related Metabolic Diseases. Frontiers in Endocrinology. 2020;11.

18. Bookout AL, De Groot MH, Owen BM, Lee S, Gautron L, Lawrence HL, et al. FGF21 regulates metabolism and circadian behavior by acting on the nervous system. Nature medicine. 2013;19(9):1147–52.

19. Kim AM, Somayaji VR, Dong JQ, Rolph TP, Weng Y, Chabot JR, et al. Once-weekly administration of a long-acting fibroblast growth factor 21 analogue modulates lipids, bone turnover markers, blood pressure and body weight differently in obese people with hypertriglyceridaemia and in non-human primates. Diabetes, Obesity and Metabolism. 2017;19(12):1762–72.

20. Marjot T, Moolla A, Cobbold JF, Hodson L, Tomlinson JW. Nonalcoholic Fatty Liver Disease in Adults: Current Concepts in Etiology, Outcomes, and Management. Endocrine Reviews. 2019;41(1):66–117.

21. Younossi ZM, Koenig AB, Abdelatif D, Fazel Y, Henry L, Wymer M. Global epidemiology of nonalcoholic fatty liver disease-Meta-analytic assessment of prevalence, incidence, and outcomes. Hepatology. 2016;64(1):73–84.

22. Challa TD, Wueest S, Lucchini FC, Dedual M, Modica S, Borsigova M, et al. Liver ASK1 protects from non-alcoholic fatty liver disease and fibrosis. EMBO Molecular Medicine. 2019;11(10):e10124.

23. Sakauchi C, Wakatsuki H, Ichijo H, Hattori K. Pleiotropic properties of ASK1. Biochimica et Biophysica Acta (BBA) - General Subjects. 2017;1861(1, Part A):3030–8.

24. Shiizaki S, Naguro I, Ichijo H. Activation mechanisms of ASK1 in response to various stresses and its significance in intracellular signaling. Adv Biol Regul. 2013;53(1):135–44.

25. Ichijo H, Nishida E, Irie K, ten Dijke P, Saitoh M, Moriguchi T, et al. Induction of apoptosis by ASK1, a mammalian MAPKKK that activates SAPK/JNK and p38 signaling pathways. Science. 1997;275(5296):90–4.

26. Keenan AB, Torre D, Lachmann A, Leong AK, Wojciechowicz ML, Utti V, et al. ChEA3: transcription factor enrichment analysis by orthogonal omics integration. Nucleic Acids Research. 2019;47(W1):W212–W24.

27. Jiang Q, Li F, Shi K, Wu P, An J, Yang Y, et al. ATF4 activation by the p38MAPK-eIF4E axis mediates apoptosis and autophagy induced by selenite in Jurkat cells. FEBS Lett. 2013;587(15):2420–9.

28. Schaap FG, Kremer AE, Lamers WH, Jansen PLM, Gaemers IC. Fibroblast growth factor 21 is induced by endoplasmic reticulum stress. Biochimie. 2013;95(4):692–9.

29. Wan X-s, Lu X-h, Xiao Y-c, Lin Y, Zhu H, Ding T, et al. ATF4- and CHOP-Dependent Induction of FGF21 through Endoplasmic Reticulum Stress. BioMed Research International. 2014;2014:807874.

30. Nikolic I, Leiva M, Sabio G. The role of stress kinases in metabolic disease. Nat Rev Endocrinol. 2020;16(12):697–716.

31. Erickson A, Moreau R. The regulation of FGF21 gene expression by metabolic factors and nutrients. Horm Mol Biol Clin Investig. 2016;30(1).

32. Pitale PM, Gorbatyuk O, Gorbatyuk M. Neurodegeneration: Keeping ATF4 on a Tight Leash. Frontiers in Cellular Neuroscience. 2017;11:410.

33. Wortel IMN, van der Meer LT, Kilberg MS, van Leeuwen FN. Surviving Stress: Modulation of ATF4-Mediated Stress Responses in Normal and Malignant Cells. Trends Endocrinol Metab. 2017;28(11):794–806.

34. Wu D, Liang J. Activating transcription factor 4: a regulator of stress response in human cancers. Frontiers in Cell and Developmental Biology. 2024;12.

35. Tobiume K, Matsuzawa A, Takahashi T, Nishitoh H, Morita Ki, Takeda K, et al. ASK1 is required for sustained activations of JNK/p38 MAP kinases and apoptosis. EMBO reports. 2001;2(3):222–8-8.

36. Berglund ED, Li CY, Bina HA, Lynes SE, Michael MD, Shanafelt AB, et al. Fibroblast growth factor 21 controls glycemia via regulation of hepatic glucose flux and insulin sensitivity. Endocrinology. 2009;150(9):4084–93.

37. Samms RJ, Smith DP, Cheng CC, Antonellis PP, Perfield JW, 2nd, Kharitonenkov A, et al. Discrete Aspects of FGF21 In Vivo Pharmacology Do Not Require UCP1. Cell Rep. 2015;11(7):991–9.

38. Véniant MM, Sivits G, Helmering J, Komorowski R, Lee J, Fan W, et al. Pharmacologic Effects of FGF21 Are Independent of the “Browning” of White Adipose Tissue. Cell Metab. 2015;21(5):731–8.

39. Kwon MM, O’Dwyer SM, Baker RK, Covey SD, Kieffer TJ. FGF21-Mediated Improvements in Glucose Clearance Require Uncoupling Protein 1. Cell Rep. 2015;13(8):1521–7.

40. Xiang M, Wang PX, Wang AB, Zhang XJ, Zhang Y, Zhang P, et al. Targeting hepatic TRAF1-ASK1 signaling to improve inflammation, insulin resistance, and hepatic steatosis. J Hepatol. 2016;64(6):1365–77.

41. Roopashree PG, Shetty SS, Suchetha Kumari N. Effect of medium chain fatty acid in human health and disease. Journal of Functional Foods. 2021;87:104724.

42. Zhou H, Urso CJ, Jadeja V. Saturated Fatty Acids in Obesity-Associated Inflammation. J Inflamm Res. 2020;13:1–14.

43. Hayakawa R, Hayakawa T, Takeda K, Ichijo H. Therapeutic targets in the ASK1-dependent stress signaling pathways. Proc Jpn Acad Ser B Phys Biol Sci. 2012;88(8):434–53.

44. Ogawa M, Kawarazaki Y, Fujita Y, Naguro I, Ichijo H. FGF21 Induced by the ASK1-p38 Pathway Promotes Mechanical Cell Competition by Attracting Cells. Curr Biol. 2021;31(5):1048–57.e5.

45. Geng L, Lam KSL, Xu A. The therapeutic potential of FGF21 in metabolic diseases: from bench to clinic. Nature Reviews Endocrinology. 2020;16(11):654–67.

46. Chui ZSW, Shen Q, Xu A. Current status and future perspectives of FGF21 analogues in clinical trials. Trends in Endocrinology & Metabolism. 2024;35(5):371–84.

47. Talukdar S, Zhou Y, Li D, Rossulek M, Dong J, Somayaji V, et al. A Long-Acting FGF21 Molecule, PF-05231023, Decreases Body Weight and Improves Lipid Profile in Non-human Primates and Type 2 Diabetic Subjects. Cell Metab. 2016;23(3):427–40.

48. Harrison SA, Rolph T, Knott M, Dubourg J. FGF21 agonists: An emerging therapeutic for metabolic dysfunction-associated steatohepatitis and beyond. Journal of Hepatology. 2024;81(3):562–76.

## References

Challa, Tenagne D., Stephan Wueest, Fabrizio C. Lucchini, Mara Dedual, Salvatore Modica, Marcela Borsigova, Christian Wolfrum, Matthias Blüher, and Daniel Konrad. 2019. ‘Liver ASK1 protects from non-alcoholic fatty liver disease and fibrosis’, EMBO Molecular Medicine, 11: e10124.

Jung, Y., M. Zhao, and K. J. Svensson. 2020. ‘Isolation, culture, and functional analysis of hepatocytes from mice with fatty liver disease’, STAR Protoc, 1: 100222.

Keenan, Alexandra B., Denis Torre, Alexander Lachmann, Ariel K. Leong, Megan L. Wojciechowicz, Vivian Utti, Kathleen M. Jagodnik, Eryk Kropiwnicki, Zichen Wang, and Avi Ma’ayan. 2019. ‘ChEA3: transcription factor enrichment analysis by orthogonal omics integration’, Nucleic Acids Research, 47: W212–W24.

Pfaffl, M. W. 2001. ‘A new mathematical model for relative quantification in real-time RT-PCR’, Nucleic Acids Res, 29: e45.

